# Residue-level predictions of the protein-protein interactions of the hepatitis B virus core and envelope proteins

**DOI:** 10.64898/2026.09.07.749370

**Authors:** Carolanne van Belleghem, Sara Rescalli, Mathilde Briday, Christophe Combet, Mario Enrique Cano Contreras, Lauriane Lecoq, Marie-Laure Fogeron, Alessandra Carbone, Anja Böckmann

## Abstract

We here predicted the interactions between the capsid (Cp) and envelope proteins (S/M/LHBs) of the hepatitis B virus using a recently established mutation-driven deep-learning model, as well as coevolution signatures that serve as markers of physical interactions and/or functional relationships. The sequence-based analyses reveal putative protein-protein interaction (PPI) hotspots in proteins, and identify abundant coevolved residues within and across proteins. We analyze the results with a focus on the intermolecular interactions between Cp and the large envelope protein LHBs, especially its disordered preS domain. We compare the predicted PPI interface sites to previous evidence on PPIs, derived from mutational analyses described in the literature. We equally integrate experimental NMR data that provide a rationale for the previous observation that spike-binding peptides inhibit core-envelope interactions. Our work sheds new light on the molecular mechanisms at play on HBV envelopment, and provides starting points for the experimental investigation of these interactions using structural and molecular virology approaches.

## Introduction

Chronic hepatitis B infection affects more than 250 million people today [1] and is the leading cause for terminal liver disease. The hepatitis B virus (reviewed in Seeger et al. [2]) is a small enveloped virus, whose genomic information codes only for a few proteins: the three envelope proteins HBs, the core protein (Cp), the polymerase P and HBx [3]. The capsid is formed by the core protein, whose different functions are driven by phosphorylation/dephosphorylation of its C-terminal part [4–8]. Cp is a 183-residue protein with two domains: the assembly domain (residues 1-140) that forms the contiguous capsid shell, and the C-terminal domain (CTD, residues 150-183) that is amongst other functions responsible for RNA packaging. The two domains are connected by a linker (residues 141-149). In infected cells, the core proteins auto assemble into capsids and concomitantly pack the pregenomic (pg) RNA, as well as a copy of the viral polymerase. Inside the capsid, the pgRNA is retrotranscribed into single-stranded (ss) DNA and then double-stranded relaxed circular (rc) DNA, generating mature capsids ready for envelopment.

The capsid structure has been investigated by a range of structural biology techniques, and besides a 3.3 Å X-ray structure [9,10] of the N-terminal assembly domain, structures of the full-length capsid have been determined by cryo-electron-microscopy (cryo-EM) [11,12], the latest to date at 2.7 Å resolution [13]. A recent manuscript describes the DNA-filled capsid [14]. The assembly domain forms mainly T=4 icosahedral capsids, made respectively from 120 Cp dimers. In the capsids, four monomers of Cp assemble in the asymmetric unit, which arrange around five-fold and quasi-sixfold axes. The core protein structure displays five α-helices, with helices 3 and 4 forming a hairpin. In the dimer, these hairpins assemble into a four-helical bundle, whose tips form the capsid spikes when assembled. Helix 5 is involved, together with the downstream loop, in interdimer interactions at the five- and quasi-sixfold capsid vertices. The highly positively charged CTD escapes as of today structure determination, even if it could be localized to regions near the inner surface of the capsid [15,16]. Also, we recently showed that it displays microsecond time scale dynamics [17]. The different Cp structures have mostly been described as similar to the initial X-ray structure, although small differences have been attributed to the absence/presence of the CTD [18], the presence of RNA as opposed to DNA [19], or due to drug binding [20].

In infectious virions, the capsid is enveloped by the three lipid-interacting surface proteins SHBs (small), MHBs (medium) and LHBs (large). LHBs has three regions: preS1 (108 to 119 residues depending on the genotype) and preS2 (55 residues), constituting together preS, and the SHBs protein (226 residues). The structure of SHBs in the context of subviral particles (SVPs) has been recently solved by cryo-EM (PDB: 7TUL [21]; 8YMJ [22]; 9IYX [23]). The SVP’s building block is a dimeric protein, with two straight helices, followed by a V-shaped helix, which lays flat at the particle surface. Part of this helix is amphipathic. Proceeding inward, the density arranges into the last helix, which is U-shaped and turns back to the surface.

It is unclear how readiness for capsid envelopment is conveyed, and Summers and Mason [24] postulated early-on that a conformational change during capsid maturation might give the signal. Alternatively, the interplay of two events has been put forward to trigger this conformational change: achievement of genome maturation inside, sensed by the phosphorylation status of the CTD and transmitted by this domain to the outside. As it has been recently reported that not only mature, rcDNA containing capsids with a defined CTD phosphorylation state are enveloped, but that a variety of subviral particles are secreted, including enveloped empty capsids [25], and also phosphorylated ones [26], (reviewed in Hu et al. [27]), it becomes clear that neither CTD phosphorylation/dephosphorylation nor nucleic acid maturation are triggering envelopment. This strengthens today the hypothesis of an untriggered, spontaneous conformational change, as also described in HBV envelope maturation [28].

Mature HBV virions are secreted after envelopment of the mature rcDNA-containing capsid through the SHBs, MHBs and LHBs envelope proteins. It has been reported in the literature that naturally-occurring mutations of Cp, notably residues 97, or in 5 and 60, could lead to immature secretion [29], low secretion [30], respectively, or on the contrary, to higher genome maturity [31], associated with genotype G. This modulation of secretion has been shown often to depend only on a single mutation, but can also arise from the addition of extra amino acids, as present for instance in genotype G [31].

The F/I97L mutation associated to an immature secretion phenotype [29] is frequently observed in the context of chronic HBV, and displays a nonselective and excessive secretion of virions containing single-stranded DNA genomes, for which the mutant core protein alone is necessary and sufficient. This is an exception to the general case where virions containing mature genomes are preferentially exported. It has been forwarded that the observed phenotype is indirectly caused by a perturbed interaction between core and envelope proteins. This is supported by the observation that a mutation in preS1 (A119F) restores the wild-type-like phenotype [32]. The structure of the F97L mutant protein was previously solved using cryo-EM [13], and compared to a newly determined WT structure. The mutant and WT EM maps are described to be highly similar, with the only difference being a local effect due to the smaller L97 residue whose sidechain is displaced, leading to an enlargement of the hydrophobic pocket in the center of the spikes. It has been suggested that this pocket is directly involved in the interaction with preS during envelopment, and that its structural details could lead to less efficient interaction with the envelope proteins, and notably preS, and as a consequence to slower envelopment of the WT capsid or the F/I97L capsid - A119F preS1 double mutant [13]. We have recently, after having evidenced binding of Triton-X100 to this pocket [33], analyzed in detail the binding of a series of derived small molecules to this pocket [34]. This revealed that the Triton-X100 aromatic moiety is essential for binding, while the size of the hydrophobic chain modulates the binding affinity, as was also confirmed by cryo-EM [35].

Two additional naturally occurring point mutations have been identified in the core protein, involving residues 5 or 60, that result in low-level secretion [30]. It was shown that the P5T mutation can compensate and rescue the I97L mutation, but the related mechanism remains undisclosed today, even if modified core-envelope interactions were forwarded as possible explanation [36]. The authors concluded that the structural details of the entire Cp molecule are important for virion secretion, as residues showing modulation of envelopment are localized in diverse regions of the protein. Other mutations have been identified by Ponsel et al. [37], namely for S17, F18, L95, K96, F122, I126, R127, N136, A137, and I139, which all allowed capsid formation. However, no envelopment of such capsids was observed, neither in empty nor in genome-containing form, as well as a reduced virion formation to undetectable levels. Beyond the hydrophobic cavity, these mutations map around the base of the spike, and they have been postulated by the authors to be “candidate sites for the interaction with envelope proteins during virion morphogenesis”.

Finally, a phenotype of higher genome maturity (i.e. a further filled-in gap in the plus-strand DNA than in the wild-type virus) has been observed for the G genotype of HBV. Genotype G is special since it shows an N-terminal 12-amino-acid extension of the core protein. The insertion enhances the genome maturity of secreted virus particles, and it was postulated that this effect possibly occurs through less efficient envelopment of core particles [31]. The structure of the capsid from genotype G has been determined at a resolution of 14 Å using cryo-EM and reveals surface features corresponding to the N-terminal insertion which may partially obscure residues on the core surface lining the hydrophobic pocket essential for normal virion secretion [38]. No detailed picture could however be obtained at this resolution.

It was found previously that the process of viral maturation, where membrane-bound surface proteins package core protein capsids, is intercepted by certain peptides comprising a LLGRMKG motif [39]. It has been shown by cryo-EM that this motif binds to the capsids at the tips of dimeric spikes [40], suggesting that the tips of the spikes act as an autonomous binding platform. While it was concluded from this work that the spike tip is involved in envelope interactions, the structure revealed however only a part of the peptide, likely the residues which bind Cp with the highest binding constant. It was unclear from the work whether residues of the bound peptide remote from the spike could block additional sites beyond the spike, also considering that the spike base is the region where most envelopment-impacting mutations are localized.

Despite these advances, the molecular basis of the interaction between the envelope and core proteins during viral particle assembly remains unclear. In this study, we combined computational and experimental approaches to investigate interactions between Cp and LHBs. Specifically, we employed mutation-driven deep-learning model (MuLAN) [41], coevolutionary analysis (iBIS2Analyzer) [42], and NMR spectroscopy. Integrating computational predictions with experimental data, we contribute three new lines of evidence addressing this long-standing problem. First, we identify residues in Cp and in both domains of LHBs predicted to participate in protein-protein interactions, and we assign these loci to existing functional knowledge, and where possible, we propose functional hypotheses for newly identified sites. Second, coevolutionary analyses across independent genotype-specific sequence datasets reveal interactions signatures that not only confirm functional residues described before, but also highlight additional candidates possibly involved in envelopment. Third, we show that spike-binding peptides actually act on Cp beyond the spike tip, thereby also modifying amino acids at the spike base that are thought to mediate interaction with the envelope proteins. Our data reveal new insight into candidate residues mediating interactions between the capsid and envelope proteins, and establish a foundation for future experimental studies into the elusive capsid-envelope interplay during HBV particle assembly.

## Results

### Capsid-envelope protein interactions revealed by mutation-driven light attention networks (MuLAN)

First, we used single-sequence data to predict interaction interfaces in the capsid and envelope proteins. To this end, we applied MuLAN [41], a deep learning method that combines light attention networks [43] with pre-trained protein language models [44] to predict mutation-induced changes in binding affinity. MuLAN’s light-attention module provides residue-level scores that can be used to identify candidate protein–protein interaction interface and functional sites.

MuLAN scores were computed for Cp, preS and SHBs. Figure 1 shows the resulting scores color coded on the Cp [10] and SHBs [22] structures, as well as on the preS sequence, for genotypes B and D (genotypes A and C are shown in Figure S1, while full MuLAN scores for Cp, preS and SHBs are provided in Figure S2-S4). In Cp, the highest-scoring residues cluster in two regions of the structure: (i) the N-terminal segment forming part of the baseplate of the two spike helices, and (ii) the region comprising amino acids in and around the hydrophobic pocket, centered near residue 100. In addition, residues within the intrinsically disordered C-terminal part (amino acids 160-170) display some of the highest scores. These high-scoring regions are largely conserved across genotypes (Figure S2).

**Figure 1.**
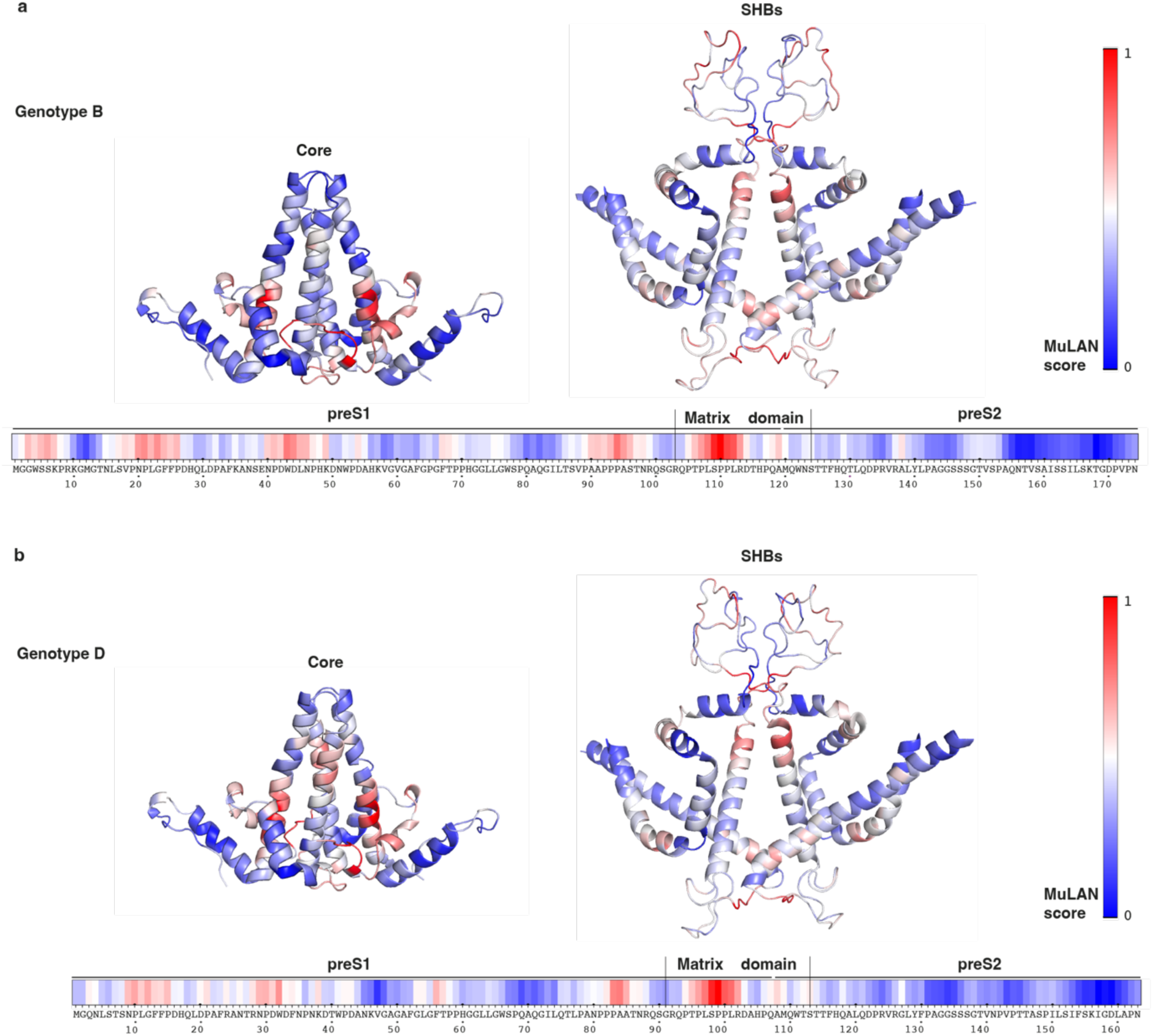
Predicted interfaces as reflected in MuLAN [41] scores for Cp, preS and SHBs. (a) Residues in the capsid (PDB: 1QGT [10]) , SHBs (PDB: 8YMJ [22]) structure and preS proteins sequence from genotype B are colored with positional scores from the MuLAN prediction (Figure S2-4). (b) Residues in the capsid (PDB: 1QGT [10]) and SHBs (PDB: 8YMJ[22]) structures, and preS protein sequence from genotype D are colored with positional scores from the MuLAN prediction (Figures S2-4).

In preS, the region with highest scores in all genotypes is the proline-rich region in the matrix domain (around amino acid 100 in genotype D and 110 in other genotypes), which harbors two Thr and Ser residues which we have previously shown to be phosphorylated on recombinant expression in a cell-free system, and also in an engineered bacterial system co-expressing the MAP kinase [45]. A second region with high scores is also a proline-rich stretch preceding the first one, harboring a PPP motif in all genotypes. A third region is formed by residues roughly in positions 20-50 (10-40 in genotype D), which include the sodium-taurocholate cotransporting polypeptide (NTCP) receptor binding site[46].

Interestingly, the very N-terminal amino-acid stretch in genotypes A-C, GP/WSSK, is also highlighted as an interaction motif. Some additional motifs displaying lower scores are equally observed (Figure S3).

In SHBs, several sites exhibit high scores, with the highest values localizing to segments of both the cytosolic (positioned at the base of the SHBs structure in Figure 1) and the antigenic loops (at the apical region). Intriguingly, consistently high scores are assigned to the hydrophobic LIFLLVLLDY motif located at the C-terminal end of helix 2, immediately preceding the antigenic loop (Figure S4). Additionally, more moderate scores are observed in the kink of the amphipathic helix 3 and within the second segment of the V-shaped helix 4.

BOLTZ-2 [47] or AlphaFold-Multimer [48] can be used to predict the three-dimensional structures of protein complexes to assess whether the predicted interfaces are consistent with the residues identified by iBIS2Analyzer and MuLAN. However, in our case, the predicted folds show poor confidence, and place preS and SHBs in biologically unlikely positions, such as preS occupying core protein interfaces, or SHBs at the bottom of the capsid. We conclude that the predictive capacities for protein complexes are still beyond the power of what presently can be successfully done.

### Coevolution analysis identifies intermolecular capsid-envelope interaction motifs

To identify residues potentially involved in capsid-envelope interactions, we analyzed coevolutionary signals between LHBs and Cp across HBV genotypes using the iBIS2Analyzer webserver, which implements the iBIS2 algorithm [42]. iBIS2 identifies pairs, or higher-order groups, of residues that show statistically significant patterns of correlated mutations in multiple sequence alignments and that are therefore likely to participate in interaction interfaces. The iBIS2Analyzer platform provides results through an intuitive web interface, along with a comprehensive textual output listing residues exhibiting coordinated mutational patterns.

Protein sequences were extracted from the HBVdb [49] and were used as input to iBIS2Analyzer (Supplementary Data Files 1-8). We filtered the resulting output (given in Supplementary Data Files 9-16) to exclusively analyze intermolecular coevolution clusters, which were only observed in genotypes A to D, but not in E to H, where fewer sequences were available. We thus further inspected coevolution clusters for genotypes A-D respectively. We concentrated on those showing a P-value score smaller than or equal to 10^-10^. We also applied a selection based on the type of amino acids in the mutation pattern, and retained only those which were non-conservative, i.e. induced a change in chemical properties. Figure 2 shows the results for genotypes B and D, for which most clusters were identified. The positions are shown on the structures of Cp (PDB: 1QGT [10]) and SHBs (PDB: 8YMJ [22]). For preS, which is intrinsically disordered [45], coevolving residues are instead shown along the sequence. The preS1 and preS2 segments are also indicated in the sequence, together with the matrix domain located at their interface.

**Figure 2.**
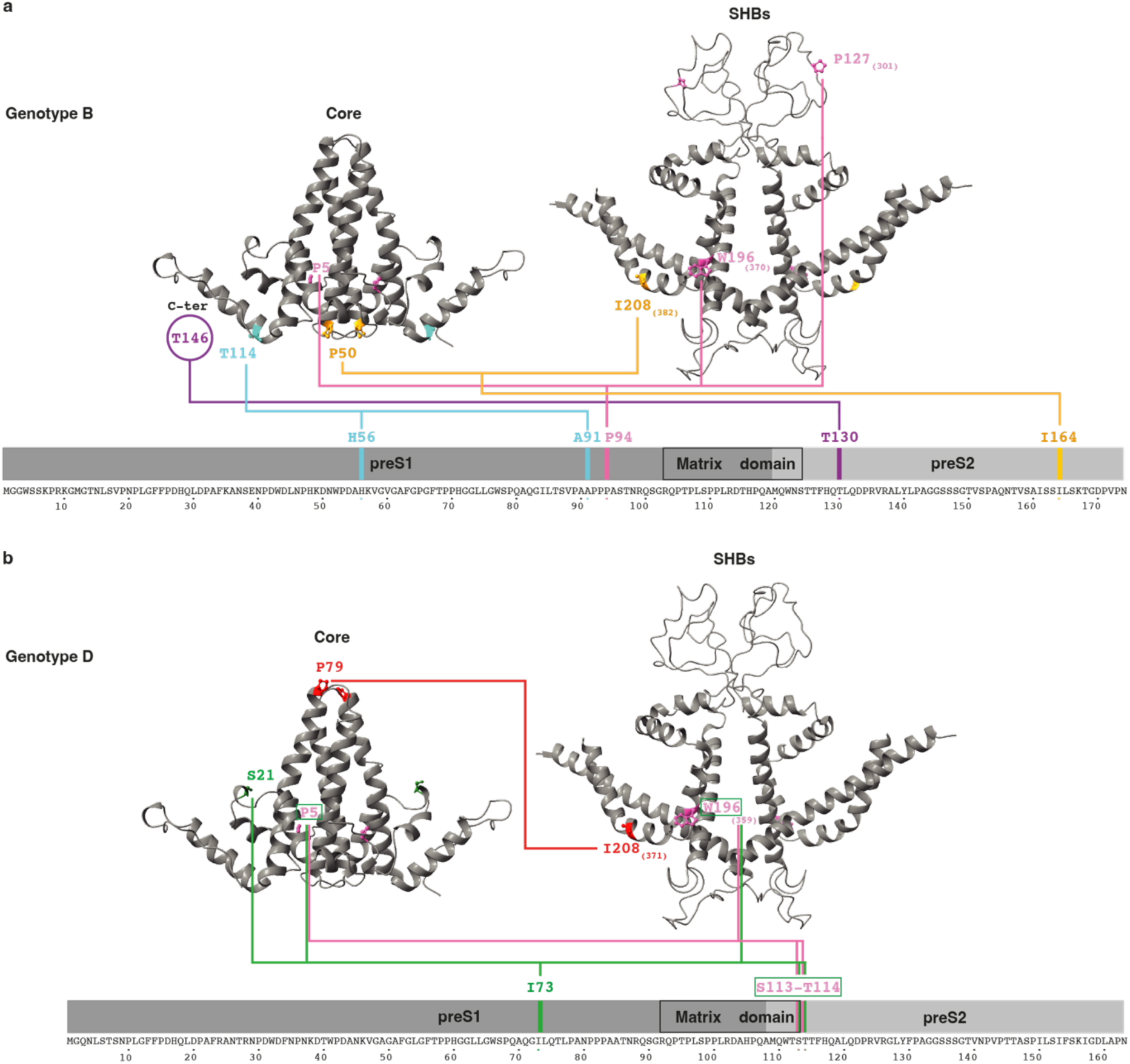
Coevolution clusters detected in the Cp dimer, preS and SHBs. (a) Genotype B shows four coevolution motifs shown in purple, blue, yellow and pink. Amino-acid numbers on the SHBs structure are given for SHBs numbering and in brackets for LHBs. (b) Genotype D shows four coevolution motifs, represented in pink, red and green. Residues Cp P5, preS2 S113/T114 and SHBs W196 are detected in both green and pink clusters. The residue positions of each cluster are connected by lines. Structures used in the plots are from Cp (PDB: 1QGT [10]), SHBs (PDB: 8YMJ [22]).

In genotype B (Figure 2a), iBIS2Analyzer identified four distinct motifs of coevolving residues. Two of them show coevolving positions in each of the three proteins. The first corresponds to coevolving residues Cp P5, preS1 P94 and SHBs W196/P127 (for equivalent sequence positions in LHBs see Supplementary Data File 17). The second shows that residue Cp P50 coevolves with preS2 I164 and SHBs I208. The two other motifs reveal a coevolution pattern exclusively between Cp and preS. The first shows that Cp T146 coevolves with preS2 T130. The second motif involves Cp T114, which is coevolving with preS1 H56 and A91.

In genotype D (Figure 2b), we identified three distinct motifs of coevolving residues. Two of them show coevolution between Cp, preS and SHBs. Firstly, again Cp P5 is detected to coevolve, this time with preS2 S113/T114 of the matrix domain, and with SHBs W196. Second, Cp P5/S21 coevolve with preS1 I73; preS2 S113/T114 at the end of the matrix domain; and SHBs W196. The third motif involves residue Cp P79, localized in the spike, and SHBs I208.

For genotypes A and C (Figure S5), 2 and 1 motifs of coevolving residues were identified respectively. In genotype A, one motif indicates coevolution of Cp E77 in the spike region with preS2 Q132, and SHBs D144/G145/F220. The second motif shows coevolution between Cp I116 and SHBs V47. In genotype C, Cp P20 is identified to coevolve with preS2 Q129.

### Solid-state NMR reveals the full effect that inhibitory peptide binding has on Cp

It has been previously shown that the two peptides MHRSLLGRMKGA and GSLLGRMKGA (referred to in the following as Oct1 and Oct2, respectively) bind to Cp and can inhibit virion formation [39]. The use of peptide microarrays revealed that it is the SLLGRM motif that binds at the spike tip that could correspond to the 5-6 residues seen in the cryo-EM density [40]. Solution NMR studies were conducted beforehand on Oct1/2 bound to the Cp dimer [50], and revealed that the peptide caused ^1^H and ^15^N chemical shift perturbations (CSPs), including in residues beyond the spike tip. In this study however, signals of the spike tip itself could not be detected, since their dynamic behavior make them disappear in the Cp dimer solution NMR spectra, together with several other flexible loops [50].

In order to further investigate the binding, we recorded solid-state NMR spectra of the two peptides bound to the assembled Cp, in order to analyze their impact on all residues, including also on side-chain chemical shifts. Beforehand, we confirmed, using isothermal titration calorimetry, that both peptides bound to Cp (Figure S6). Determined affinities (5-6 μM) compare within an order of magnitude to those described the literature, where 2 and 68 μM affinity was reported [40,50].

Figure 3a,b shows the 2D DARR spectra recorded on Cp in presence of Oct1 or Oct2. The resonance assignments could be transferred from the apo form reported in reference [33]. Figure 3c,d shows the derived CSPs, which display larger maxima for Oct2. The CSPs have been reported on the Cp structure (PDB: 1QGT [10]) in Figure 3e,f. When comparing the CSPs induced by Oct1 and Oct2, one can see that both peptides induce perturbations in similar regions of the Cp structure. The residues impacted by the binding of the peptides localize to the spike, but also further down the helices. It is clear that CSPs are significant in the very N-terminal residues, and in those between residues 55-100, with the changes at the spike tips not much larger than in the remaining regions. The affected residues go clearly beyond the ones reported in the solution NMR study, and add residues in the N-terminal part to the affected amino-acids, whereas this part was described as unaffected before [50].

**Figure 3.**
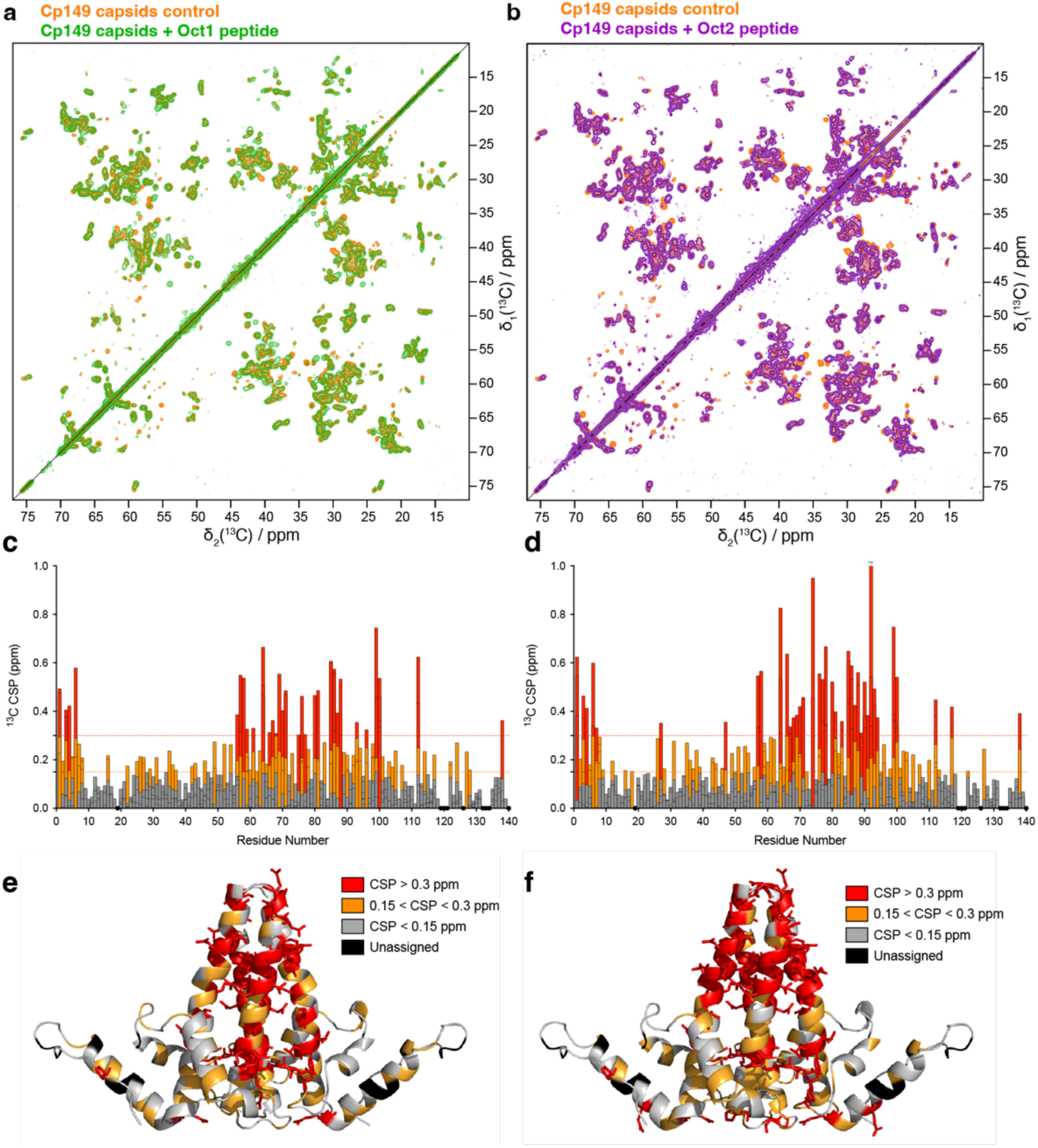
Binding of Oct peptides to Cp as seen by NMR. (a) Comparison of the aliphatic region of 2D DARR spectra of labelled ^13^C^15^N-Cp149 reassembled capsids in absence (orange spectrum) or in presence of Oct1 (green spectrum); (b) Comparison of the aliphatic region of the 2D DARR spectra of labelled ^13^C^15^N-Cp149 reassembled capsids in absence (orange spectrum) or in presence Oct2 (purple spectrum). Experimental parameters are detailed in Table S1. (c,d) Residues of Cp149 dimer impacted by Oct1 and Oct2 binding respectively. CSPs graphs show, for each residue, the difference of chemical shift between the unbound and the bound form, with every bar representing a single resonance. Bars are colored in grey, orange and red for small, medium and large CSPs, respectively. Black circles represent unassigned residues. (e) and (f) report the CSPs on the structure of the dimer (PDB: 1QGT [10]) for Oct1 and Oct2 respectively, mapping amino acids with small (grey), medium (orange) and large (red) CSPs. Black residues represent unassigned residues.

## Discussion

MuLAN[41] and iBIS2Analyzer [42] are two independent and complementary sequence-based approaches for predicting amino-acid residues involved in protein-protein interactions. Importantly, they rely on different types of information: MuLAN extracts residue-level interaction signals using a mutation-driven light-attention architecture trained on binding affinity changes, whereas iBIS2Analyzer detects coevolutionary signals between protein pairs from sequence alignments. We therefore use them together as independent sources of evidence, comparing their predictions to give additional weight to residues supported by convergent signals. This combined sequence-based strategy is particularly important in the present context, as it enables inclusion of the preS domain, which does not adopt a stable folded structure in isolation and remains poorly resolved by current structure prediction methods. Applying these two methods, we identified a number of residues in the Cp, preS and SHBs proteins that may contribute to PPI. We discuss in the following these different sites and confront them with previously reported functional data for each of the three proteins.

### Protein-protein interactions in the core protein

We first analyzed the capsid protein (Figure 4), and discuss in the following the results across its different regions.

**Figure 4:**
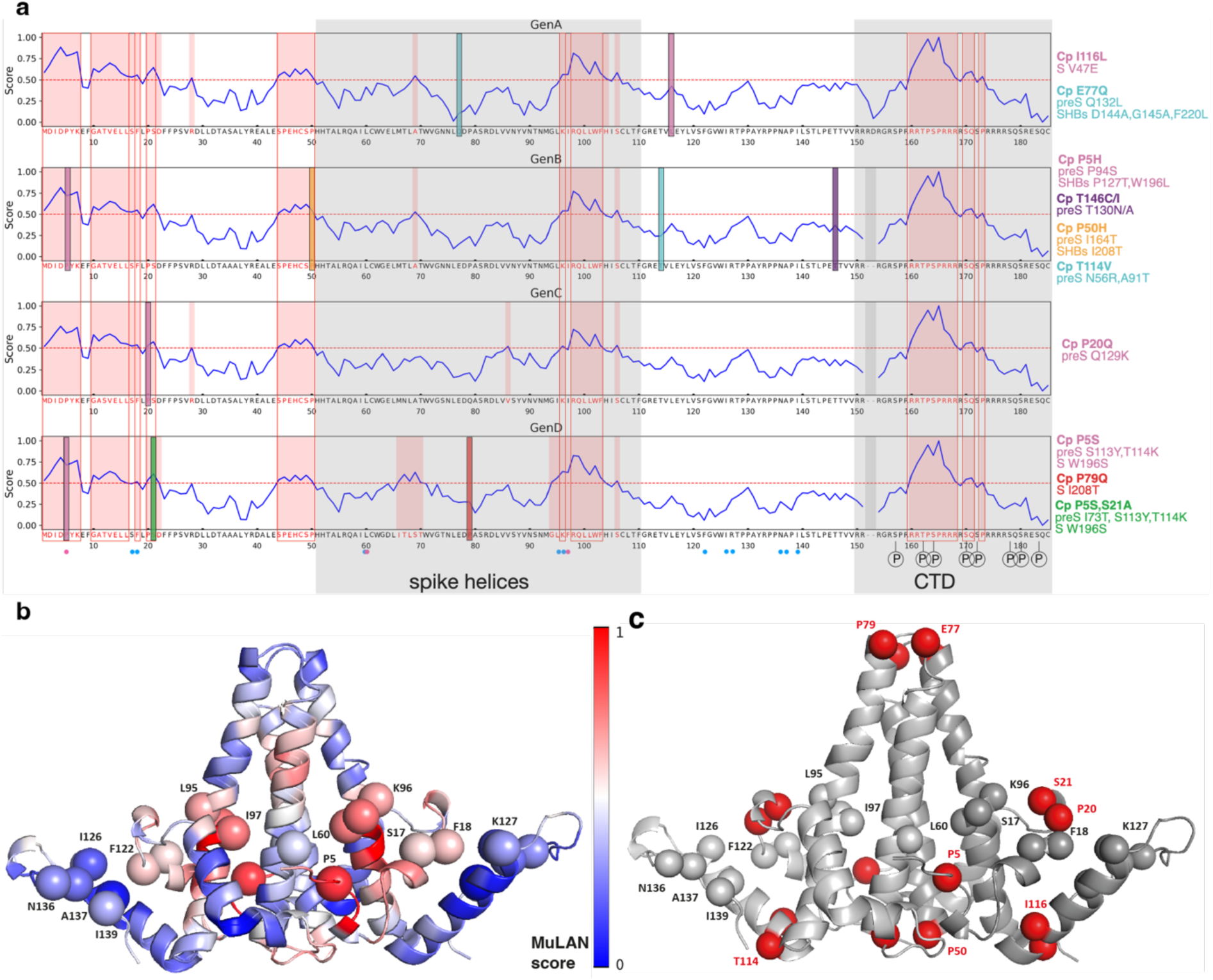
Summary of prediction results for Cp and comparison to previous knowledge. (a) Shown are scores from MuLAN predictions for all genotypes, as well as coevolving residues detected by iBIS2Analyzer. Major structural features, as the spike helices and the disordered CTD are indicated. Coevolution events are displayed on the sequence as bars, with the same colors as in the result section. The coevolving residues are given for each genotype on the right of the MuLAN scores. S or T residues that have been previously shown to be phosphorylated [56] are indicated. Mutations identified as abolishing envelopment are given by blue dots below the sequence [37], and mutations that modify secretion phenotypes [30] are indicated by pink dots. (b) The aforementioned mutations are highlighted on the structure (PDB: 1QGT [10]), where MuLAN scores are color coded on the backbone and Cα atoms of residues of interest. (c) The same residues are highlighted, together with the coevolution residues found by iBIS2Analyzer colored in red.

#### N-terminal domain

A first region that consistently shows high MuLAN scores is the N-terminal domain, with residues 1-7, 10-16, 18 and 20/21 making consensus between the different genotypes. Many of these residues form part of the spike base plate, which has been proposed as an interacting region with preS [37]. In this region, P5 emerges as a central residue: a naturally occurring P5T mutation is characterized by reduced virion secretion [30]. Also, we identified P5 to coevolve in two genotypes with SHBs W196, further revealing their connection with SHBs P127, and preS P94/S113/T114. There is no prior indication in the literature about a connection between these residues, and its functional importance remains to be established. Two other non-natural mutations, at residues S17 and F18, have been shown to abolish envelopment [37]. Both positions display significant MuLAN scores, although no coevolution signals were detected. Interestingly, two nearby residues, P20 and S21, also exhibit significant MuLAN scores, and in addition, show coevolution signals with preS and SHBs. They are predicted to coevolve with preS Q129 in genotype C, with preS I73/S113/T114 in genotype D, and with SHBs W196.

#### Loop residues lining the capsid interior around amino acid 50

Another region of interest, consistently associated with significant scores, lies immediately before residue P50. This region represents a loop pointing towards the capsid interior, just before the spike helices. A putative zinc-binding site involving Cp H47 has been proposed previously [51], and a disulfide bond between Cp C48 and Cp C183 has been described to play a role in viral particle assembly [52]. A coevolution motif is observed for Cp P50, linking this residue to preS I164 and SHBs I208. Surprisingly, SHBs I208 is also connected to the spike tip in genotype D, and to Cp P79. This suggests that Cp P50 and the spike tip are connected through their shared association with SHBs I208; however, the structural basis and functional significance of this interaction remain to be elucidated.

#### Capsid spike tip

The spike tip has been hypothesized as playing a central role in the capsid-envelope interactions, mainly supported by the finding that viral maturation is intercepted by certain peptides [39] binding to the capsids at the tips of dimeric spikes [40]. However, MuLAN scores in the spike tip are consistently low over all genotypes. In line with this, a deletion of A80 at the tip of the spike had been shown to behave as wild type with respect to virion formation, indicating that this site is not crucial for Cp-envelope interactions during particle formation [53,54]. Also, other mutations at the tip or stem of the spike had no impact on envelopment [37]. These authors therefore concluded that the inhibitory effect of the peptide is mediated by steric hindrance rather than by the blocking of a specific envelope–capsid interaction. The NMR-experimental results strongly support this conclusion, as the impact on the Cp chemical shifts of the peptide binders clearly extends beyond the spike tip, inducing significant modifications, including at the very N-terminal, around hydrophobic pocket residues and amino-acids at the spike base. These modifications can therefore be held accountable for blocking the interaction on envelopment.

Nevertheless, as mentioned above, Cp P79 in the spike does show a coevolution signal with SHBs I208. Another residue in the Cp spike tip, E77, displays a coevolution signal in genotype A, this time with preS Q132 and SHBs D144/G145 and F220. As also the case for Cp P79, this connection is in need of a rationale.

#### Hydrophobic pocket

A region that is detected to be particularly important by MuLAN localizes around residue Cp residue 100. It indeed comprises many residues lining the hydrophobic pocket [30,33,55]. Consensus for significant or high MuLAN scores are residues Cp K96 and R98 to F103, with F97 just below the threshold in genotype C. The hydrophobic pocket’s role has been the focus of much research, given its identification as a hotspot in envelopment, primarily hypothesized through the naturally occurring F97L mutation, which results in a premature envelopment phenotype [30]. F97 is localized at the bottom of the hydrophobic pocket, and its conformation is impacted by hydrophobic-pocket binding molecules [33–35]. Two residues forming part of the hydrophobic pocket, L95 and K96, shown in Figure 4a as blue dots below the sequence, have been shown to abolish envelopment when replaced by alanine [37]. The most likely interpretation for the observation of altered secretion phenotypes forwarded by the authors was a modified heterologous core-envelope protein interaction through the mutations. K96 is highlighted by MuLAN with significant scores (Figure 4a,b). Besides, three coevolution signals are identified for residues T114, I116 and T146. All three localize to the Cp base, where the Cp CTD is localized. They show correlations to preS H56/A91/T130 and V47 in the SHBs cytosolic loop (Figure 2 and S5). These interactions cannot be rationalized with current knowledge, and will be of interest as focus in further experimental approaches.

#### C-terminal domain

The last segment with high MuLAN scores is found within the C-terminal domain, with consensus on residues 160-168, 170/171 and 173. It can be seen in Figure 4a that this region contains several phosphorylated residues, as described before [56].

We also compared the results from MuLAN and iBIS2Analyzer with the mutations identified before [37,54] on the Cp structure. Figure 4b shows the MuLAN scores color coded on the structure (PDB: 1QGT[10]), where mutations that modified the envelopment phenotypes are shown as spheres. One can see that the residues localized at the spike base are characterized by high MuLAN scores, and that also the residues closer to the interdimer contacts in the capsid show significant MuLAN scores. Eight of the mutations that impact envelopment show also coevolution signals (Figure 4c), highlighting their involvement in Cp-HBs interactions.

### Protein-protein interactions in the preS domain of LHBs

We can use a similar approach to relate our predictions to the interactions of the preS domain that have been described in the literature using experimental procedures (Figure 5).

**Figure 5:**
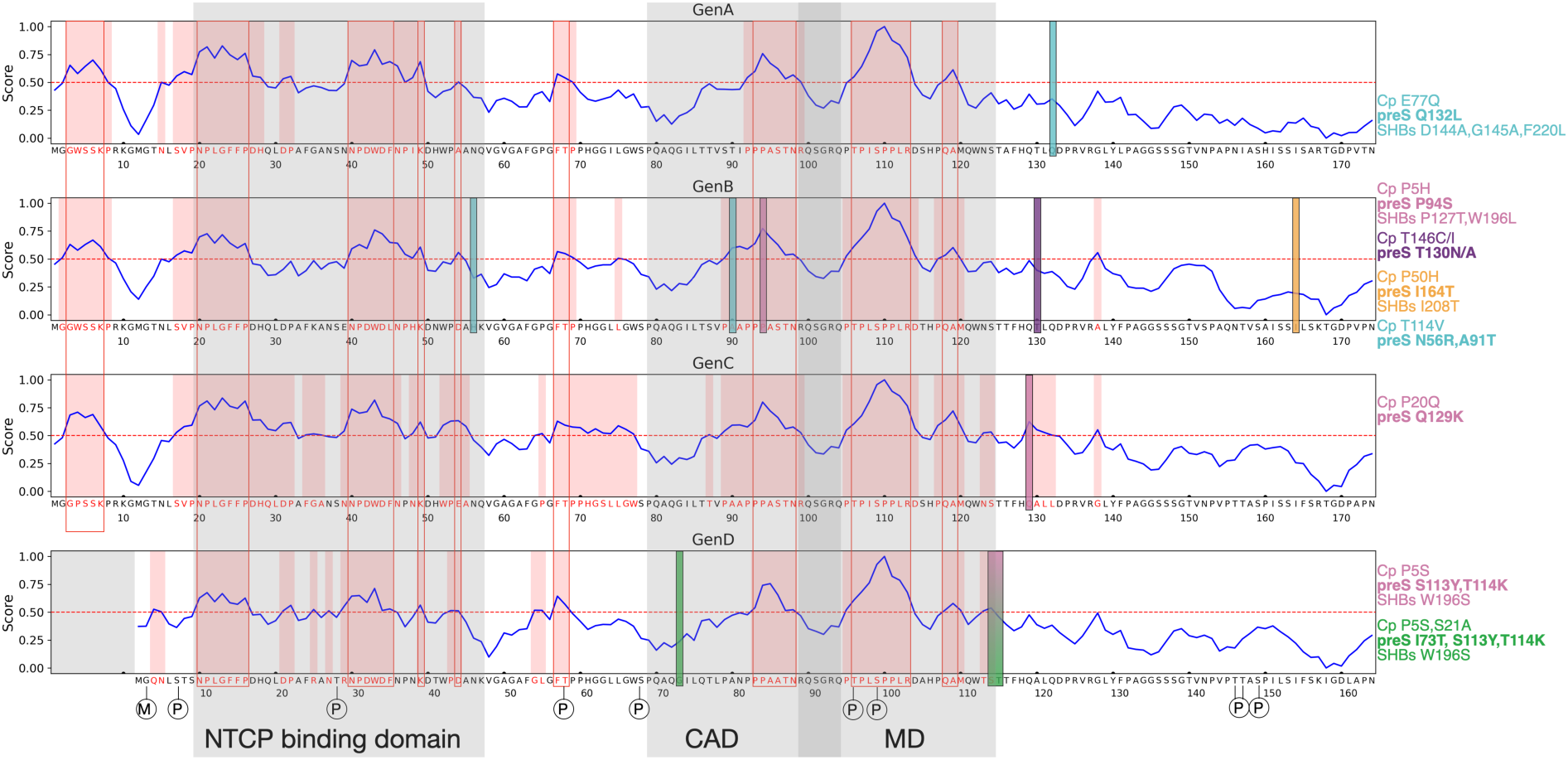
Summary of prediction results for preS and comparison to previous knowledge. Shown are scores from MuLAN predictions for all genotypes, as well as coevolving residues detected by iBIS2Analyzer. Major functional features are indicated. Coevolution events are displayed on the sequence as bars, with the same colors as in the result section. The coevolving residues are given for each genotype on the right of the MuLAN scores. S or T residues that have been shown to be phosphorylated [45] or G that is myristoylated [65] are indicated by P and M respectively. Major features, as the NTCP binding domain, the cytosolic anchorage determinant (CAD) and the matrix domain (MD) are indicated.

#### The N-terminal preS residues present in genotypes A-C

Interestingly, the very N-terminal residues in preS from genotypes A-C are consistently highlighted by significant MuLAN scores. The SSK motif represents a typical (S/T)-X-(R/K) protein kinase C (PKC) phosphorylation site, and may be used as such in genotype A-C preS. Since phosphorylation patterns were only established for genotype D [45], it is unclear if this Ser residue is phosphorylated in vitro. While an active protein kinase C fragment (named PKM) was shown to be present in the HBV core particle [57], no interaction of PKC with LHBs has been described to our knowledge.

#### The preS sodium-taurocholate cotransporting polypeptide (NTCP) receptor binding site

A major interaction partner of preS is the HBV receptor (NTCP [46]), which has been shown to interact mainly with preS residues up to position 48 (genotype D) [58]. Within this region, the conserved sequence NPLGFFPDHQ is particularly important for binding to NTCP [46], and it is for residues within this peptide that MuLAN shows consistently high scores, together with the DWD motif towards the end of the NTCP-interaction region. At the end of the NTCP binding domain, preS H56 was identified as coevolving with Cp T114, but does not show a significant score in MuLAN. Still, adjacent to the NTCP binding site, an FT pair shows significant MuLAN scores. It comprises T57, which is phosphorylated *in vitro* [45]. This is also the case for S68, but which does not show significant MuLAN scores. To our best knowledge, no interaction partners have been described in the literature for this segment yet.

#### Cytosolic anchorage determinant (CAD)

The following element consistently identified by MuLAN is a PPAxTN motif localized between residues 90 and 100 (80-90 in genotype D). This region forms part of the so-called cytosolic anchorage determinant (CAD), which is the translocation control region whose interactions allow to keep preS in the cytosol for Cp envelopment. Deletion of this sequence relieves the suppression of co-translational preS translocation, yielding uniform e-preS [59]. A likely interactor here is Hsc70 [59], and MuLAN predictions further refine the interaction site to the PPAxTN stretch. Genotype B preS A91 shows a coevolution signal with preS H56 and Cp T114, and P94 shows a coevolution signal with Cp P5 and SHBs P127/W196. The peptide containing this conserved PxxPP^94^P motif tested negative [60] with respect to its role in envelopment. However, PxxP short linear motifs are known to play important roles in linking different proteins together in signaling pathways, and the motif can, for instance, recruit kinases or other signaling molecules to specific cellular locations [61].

#### Matrix domain

Further progressing to the C-terminal of preS, the next stretch highlighted in all MuLAN predictions is the PTPxSPPLR motif, with x being L or I. This motif is at the heart of the preS matrix domain, which has been identified as the peptide stretch of central importance in envelopment [62,63]. Indeed, it has been shown that deletions in preS up to residue 102 in genotypes A-C still allowed virion morphogenesis, whereas deletion of further 8 or more amino acids destroyed this function [64]. It is thereby unlikely that the regions/residues identified both by MuLAN and iBIS2Analyzer in preS preceding position 102 point to interactions with Cp during envelopment. Also, Shih and coworkers have shown that preS A119F (genotype A) can set off, in the preS mutated form, the Cp F/I97L mutation [55]. It can be seen in the MuLAN profiles that in genotype A-C residues 118/119 (genotype D 107/108) are predicted as PPI sites. Also, a synthetic peptide corresponding to amino acids 96-116 (genotyope D) has been described as the best binder [60]; this comprises many of the residues showing high MuLAN scores in the matrix domain. The matrix domain interestingly also contains T95 and S98 residues that we previously showed to be phosphorylated *in vitro* [45]. One can highlight as well the presence of a PxxP motif, which importance in functional interactions has been described above. Surprisingly, no coevolution of matrix domain residues with Cp (besides S113, see below) have been identified; this is most likely related to the low mutation rate of the constituent residues, rather than absence of interactions. However, right after the matrix domain, preS S113/T114 (genotype D) are predicted as interacting with Cp P5 and SHBs W196 in genotype D, and furthermore a Cp P5 and SHBs W196 interaction is detected also in genotype B. preS S113/T114 form part of the peptide motif which has been shown to be relevant in envelopment [60]; preS S113 is part of the matrix domain; and P5 has been identified as key residue in envelopment. preS P94 and S113 are predicted by MuLAN to be involved in PPI, but not preS T114 and SHBs W196.

#### C-terminal part of preS

The portion of preS between the matrix domain and SHBs shows consistently poor MuLAN scores. However, there are four more coevolution signals that connect residues from this part to both the Cp and the SHBs protein. Three of them localize around residue 130, (Q129/T130/Q132), amongst which only Q129 shows a significant MuLAN score. The fourth coevolution event is detected for I164. It is worth mentioning that three further phosphorylation sites have been evidenced in this domain [45].

### Protein-protein interactions in the SHBs protein

Finally, we analyze the results of the predictions for the SHBs protein (Figure 6).

**Figure 6:**
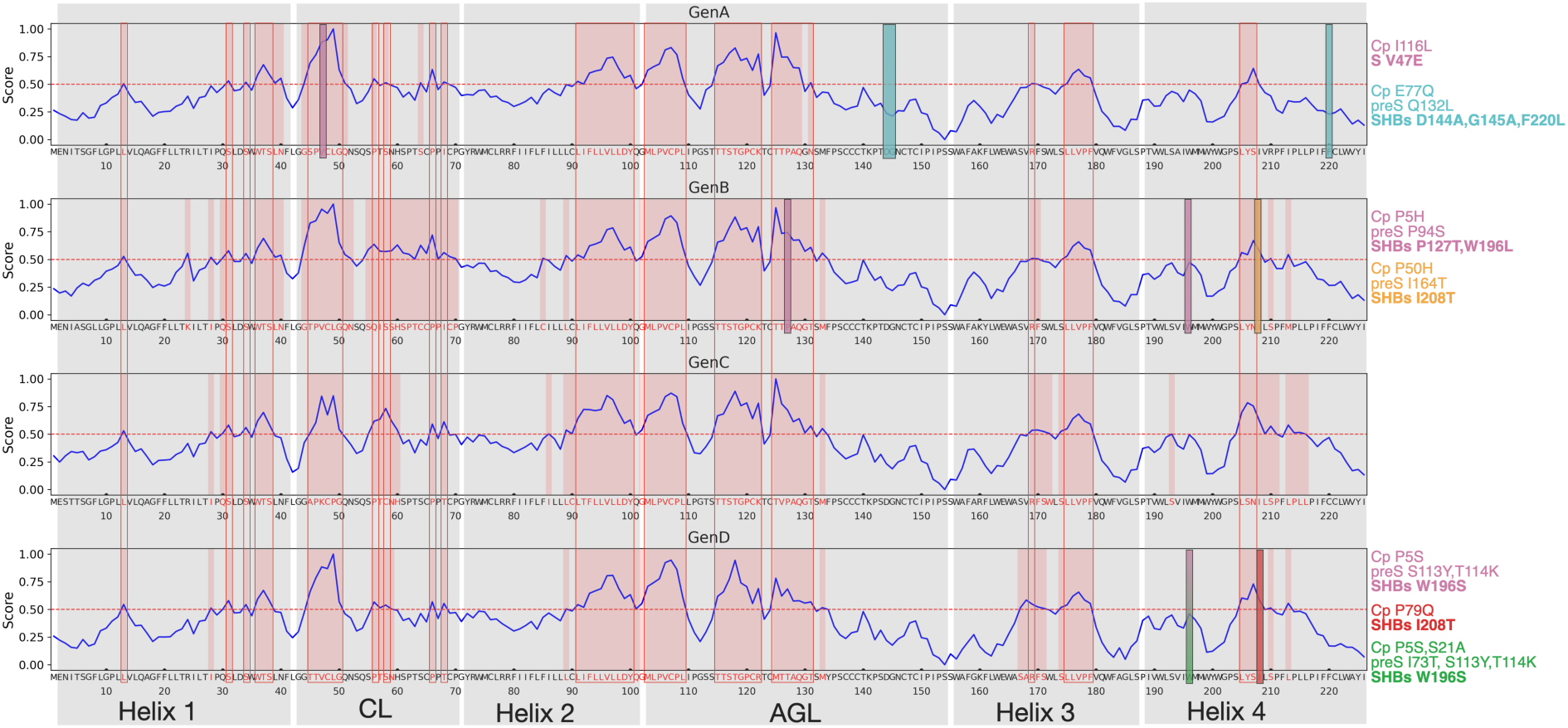
Summary of prediction results for SHBs and comparison to previous knowledge. Shown are scores from MuLAN predictions for all genotypes, as well as coevolving residues detected by iBIS2analyzer. Major structural motives are indicated. Coevolution events are displayed on the sequence as bars, with the same colors as in the result section. The coevolving residues are given for each genotype on the right of the MuLAN scores.

#### Helix 1 (H1) & cytosolic loop (CL)

Significant MuLAN scores light up systematically in residue L13 of H1. Further downstream H1, four polar residues (S31, S34, T37, S38), as well as a W36 display significant MuLAN scores. How the coevolution of SHBs cytosolic loop V47 with Cp I116 fits in the picture remains unclear at the moment; I116 is not displayed on the Cp surface. Still, considering the high number of sequences, and the conservative cutoff used, this unexpected residue pair might reveal a yet unknown functional interaction. Of interest, it has been previously demonstrated that CL deletion mutations SHBsΔ35–39, SHBsΔ40–46, SHBsΔ50–56, and SHBsΔ56–59 blocked virion production [66]. Point mutations however did not have this effect [67].

#### Helix 2 (H2)

H2 is a transmembrane helix that spans from the cytosol to the ER. On the cytosolic part, it is limited by the cytosolic loop, and on the ER lumen by the antigenic loop. The C-terminal end of this helix displays high MuLAN scores, which is rather unexpected since it seems hidden, likely covered up by lipid molecules. However, this part actually interacts in the SHBs tetramer structure with the neighboring C-terminal part of the U-shaped helix 4 [21]. This interaction likely does not exist as such for LHBs at the start of envelopment, where the C-terminal of helix 4 is likely to point more towards the cytosol, to allow an i-preS conformation [59]. Interaction of cellular factors with SHBs could further sustain an i-preS orientation. Interestingly, L14 in H1 consistently shows significant MuLAN scores; this residue is involved in intermolecular H1 interactions that are likely to be untied in the i-preS form.

#### The antigenic loop (AGL)

The AGL is involved in the initial attachment of HBV to the host cell, likely through interactions with heparan sulfate proteoglycans (HSPGs) [68]. This facilitates binding to the primary HBV receptor, the NTCP. The AGL is a highly complex structure containing eight Cys and five Pro residues. It mediates attachment to immobilized heparin and to differentiated HepaRG cells [68]. And indeed, MuLAN scores are high in the surface exposed N-terminal part of the AGL; the C-terminal portion hidden inside the loop structure shows low scores throughout all genotypes. While the two stretches ^105^PVCPL and ^118^TGPCR (genotype D) which show high MuLAN scores have been shown to be important for infectivity [69], further residues, localizing around the remaining cysteine residues of the AGL (^137^CCCTKP and ^146^NCTCI), are not highlighted by MuLAN, but are important for both heparin binding and infectivity [69]. The lack of predicted interactions for these regions suggests that their role is more likely to maintain the specific disulfide-bonding pattern, rather than to engage in direct interactions. Two coevolution events are detected in the AGL. First, genotype A SHBs D144/G145, not predicted by MuLAN to be involved in an interaction, pairs up with Cp E77 and preS Q132. While an interaction with E77, located in the Cp spike tip, is difficult to imagine with the AGL, e-preS could interact with the AGL, possibly influencing virion morphology and infectivity. Indeed, SHBs D144/G145 are important neither to heparin interactions, nor infectivity [69,68]. The same holds for the genotype D-equivalent position to genotype B SHBs P127, which pairs up with Cp P5, preS P94 and SHBs W196. Again, its interaction with e-preS could cause the predicted mutational interdependency. MuLAN shows consistently high scores for this residue, which is however not conserved in genotype D. Attachment of HBV and HDV particles to heparin has been shown to be dependent upon electrostatic interactions with SHBs R122 and K141 (genotype D) [69,68]; however, only the former shows high MuLAN scores.

#### Helix 3 (H3)

H3 is an amphipathic helix exposed on the sub-viral particle surface [21]. It shows a turn at residue R169, highlighted by MuLAN, which could be important in interactions with the lipid headgroups in which context this helix is defined. The second region with high MuLAN scores comprises a LLVPF motif, which includes a proline. This is remnant of the hydrophobic motifs that recruit chaperones and prolyl isomerases for correct folding on exit from the ribosome [70]. No coevolution event is registered for H3.

#### Helix 4 (H4)

H4 has one stretch consistently identified by MuLAN as interaction surface, the three amino acids 205-207, which are not fully conserved between the genotypes. These residues also localize nearby one of the coevolution signals detected in H4, SHBs I208 in genotypes B and D. SHBs I208 surprisingly coevolves with Cp P50 in genotype B, localized at the Cp interior and a priori not available for interaction with SHBs. Since preS I146 is part of this network, coevolution might be caused by Cp and SHBs both interacting with preS I146, which is however somewhat unexpected. In genotype D, the spike tip residue P79 is identified as a partner of SHBs I208. Another coevolution motif is detected in H4, again across two genotypes, centered on SHBs W196. In both cases, W196 pairs with Cp P5, while the associated preS residues differ, involving either P94 or S113/T114. In addition, P127 within the AGL contributes to this cluster of residues. Again, the most plausible explanation is that a preS residue evolves to interact with both Cp and the AGL during viral particle maturation. However, no known interaction accounts for these coevolution motifs in H4, which thereby likely point to undescribed interactions between these residues.

Predictive approaches currently advance at fast pace, allowing to analyze large protein sequence datasets. In this study, we investigated the protein-protein interactions involved in HBV particle formation using two independent computational methods, based on the one hand on predicted changes in binding affinity, and on the other on coevolution analysis. Because both approaches rely exclusively on sequence information, they can be applied not only to structured domains but also to intrinsically disordered regions, which are often inaccessible to structure-based methods yet play key roles in transient and dynamic interactions. These complementary sources of information provide distinct but converging signals, and their integration strengthens the confidence of our predictions. In many cases, our results reveal that the signals support the claims and data which were previously observed in the literature. But interestingly, it also points to residues that have not been previously identified as such. These residues can be yet unidentified functional hotspots that merit to be explored experimentally, as shown here at the example of the Cp spike interactions with an antiviral peptide. While the prediction of 3D structures of protein complexes is still at the edge of what is possible, harnessing efficient computational strategies using new deep learning methods allows to infer protein interactions in complex, multi partner systems, and to construct interaction landscapes for viral proteins involved in interactions essential for the viral life cycle. Application of these new methods to long-standing problems such as the HBV viral particle formation opens up new opportunities for experimental work.

## Materials and Methods

### MuLAN

MuLAN is a sequence-based deep-learning method designed to predict changes in binding affinity induced by mutations affecting protein sequences. Its light-attention module can also be applied to single sequences to assign each residue a normalized attention score in the range [0, 1], which has been shown to correlate with interacting or functional regions. MuLAN [41] is available at http://gitlab.lcqb.upmc.fr/lombardi/mulan under the CC-BY-NC-SA 4.0 License. The program was run using standard parameter sets.

### iBIS2Analyzer

iBIS2Analyzer is a web server implementing the BIS2 algorithm, which is designed to detect coevolutionary signals within and between protein sequences. The method detects groups of positions in multiple sequence alignments that show statistically significant patterns of correlated mutations. These groups of coevolving residues can provide evidence for structural or functional relationships, including potential interaction sites or conserved functional networks. The iBIS2Analyzer web server is available at https://ibis2analyzer.lcqb.upmc.fr. We used as input 1347 (genotype A), 2291 (genotype B), 3573 (genotype C), 1843 (genotype D), 460 (genotype E), 303 (genotype F), 76 (genotype G) and 34 sequences (genotype H), corresponding to the sequences available in the HBVdb (https://hbvdb.lyon.inserm.fr) [49]. Sequences were extracted from the HBVdb and concatenated in the following order: Cp (1-183), XXXXX, LHBs (190-577), aligned and saved in FASTA format accepted by iBIS2Analyzer (Supplementary Data files 1-8). X stands for a separator between the two proteins.

The coevolution clusters were ranked by their p-values from smallest to largest. The analysis of the clusters for each genotype respects a selection limit for each one, with only values below 10^-10^ being retained. Only intermolecular interactions were taken into account. In the intermolecular interaction group below the threshold limit, a selection is applied to the types of mutations observed; only mutations that induce a change in chemical properties are retained. All occurrences and subtrees are shown in the Supplementary Data Files 9-16 for the different genotypes. Supplementary Data File 17 provides equivalent sequence positions between experimentally solved 3D structure sequences (PDB:1QGT [10] and PDB:8YMJ [22] i.e. INSDC:AY057948) and the 16 HBVdb database reference sequences representing genotypes A to H, aligned by means of MUSCLE 5.3 multiple sequence alignment algorithm.

### Expression and purification of Cp

^13^C-^15^N-Cp149 capsids were expressed, purified, disassembled as Cp dimers and reassembled into capsids as described in reference [33]. The freshly prepared ^13^C-^15^N-Cp149 reassembled capsids were dialyzed in 50 mM HEPES pH 7.5, 5 mM DTT and incubated with 4 molar equivalents of Oct1 or Oct2 peptides for 2 hours at room temperature. The samples were then concentrated using Amicon Ultra centrifugal filter units (Merck, 50 kDa cut-off) to about 20 mg/ml in 1 mL and sedimented into 3.2 mm zirconium rotors by ultracentrifugation (200,000 *g*, 14 h, 4 °C) using a home-made filling tool [71]. Rotors were immediately closed after the addition of 1 µL of saturated DSS solution for chemical-shift referencing. A control sample of Cp149 capsids reassembled in absence of Oct peptides was also prepared in order to measure the chemical shift perturbations (CSPs) between the reference spectrum and those with the Oct peptides.

### NMR

2D ^13^C DARR [72] spectra were recorded using a 3.2 mm triple-resonance (^1^H, ^13^C, ^15^N) wide-bore probe head at a static magnetic field of 18.8 T corresponding to 800 MHz proton resonance frequency (Bruker Avance II) and at a 17.5 kHz MAS frequency. The ^15^N-insert was removed from the probe to increase the signal-to-noise ratio. All spectra were referenced to DSS and recorded at a sample temperature of 4 °C according to the resonance frequency of the supernatant water. Assignments of the DARR spectra were derived from those presented in reference [73] (BMRB accession number 28122) of Cp149 capsids reassembled in absence of Oct. All spectra were processed using TopSpin 4.0.3 (Bruker Biospin) and analyzed with the CcpNmr Analysis [74] package, Version 2.4.2. For Oct1 and Oct2 peptides, chemical-shift differences between Cp149 reassembled capsid without and with ligand were calculated for each carbon atom according to: Δ*δ* = *δ*[*bound*]− *δ*[*unbound*].

### ITC

Freshly prepared Cp149 reassembled capsid samples were dialyzed overnight at 4 °C into 50 mM HEPES buffer at pH 7.5. The protein concentration was determined by UV-Vis absorbance measurements using a Nanodrop. Oct peptides were dissolved in the same HEPES buffer used for the protein dialysis to ensure buffer matching between the protein and ligand solutions. Stock solutions of the peptides were prepared at 1 mM and subsequently diluted to a final concentration of 400 µM. The protein was used at a concentration of 40 µM in the ITC cell, maintaining a typical ligand-to-protein ratio of 10:1 to ensure saturation binding conditions. All ITC experiments were performed using a MicroCal iTC200 instrument (Malvern Panalytical, Malvern, Worcestershire, UK) at a constant temperature of 25°C. The titration protocol consisted of an initial injection of 1 μl of the Oct peptide solution (injection duration: 2 s), followed by 24 subsequent injections of 1.6 μl each (injection duration: 3.2 s), with a spacing of 2 minutes between injections to allow the system to return to baseline. Control experiments were performed under identical conditions and used for data correction in subsequent analysis. The raw ITC data were analyzed using the MicroCal Origin software package. Binding isotherms were fitted to a single site binding model to determine thermodynamic parameters.

## Data availability statement

All data generated or analyzed during this study are included in this published article and its supplementary information files. NMR spectra are provided on request by the authors.

## Supporting information

Sequence alignments

iBIS output

residue mappings

## Acknowledgements

Financial supports from the agence nationale de recherche sur le sida et les hépatites virales – maladies infectieuses émergentes (ANRS-MIE https://anrs.fr) (ECTZ 242442 to AB, ECTZ 243771 to CvB and ECTZ246688 to CC), from PostGenAI@Paris within the framework of ANR/France 2030 (https://anr.fr/) (ANR-23-IACL-0007 to AC), from ANR ALLEGRO (ANR-23-CE11-0006, to AC), and the Institut Universitaire de France (https://www.iufrance.fr/) (to AC) are acknowledged.

## Author contributions

A.B. and A.C. conceived and planned the work. C.v.B., S.R. and C.C. performed the bioinformatics analyses, with support from M.E.C.C., and prepared figures. M.B. and L.L. conducted the NMR experiments and related analyses. M.B and M.-L.F. performed ITC measurements. A.B. and A.C., with the help of all authors, drafted the manuscript. All authors reviewed and approved the final manuscript.

