## Supporting Information for "Residue-level predictions of the protein-protein interactions of the hepatitis B virus core and envelope proteins"

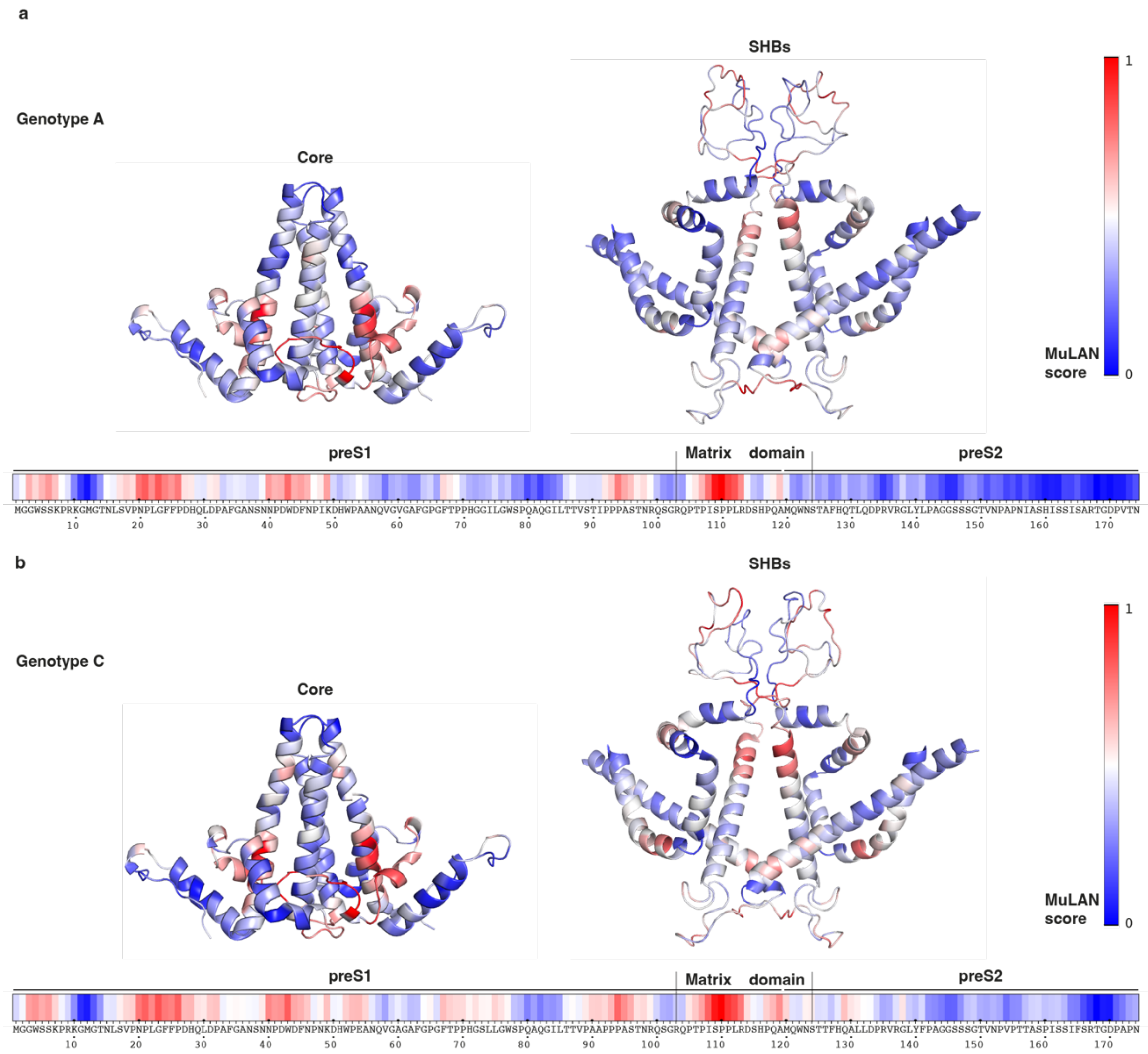

**Figure S1. Predicted interfaces as reflected in MuLAN scores for Cp, preS and SHBs.** a. Residues in the capsid (PDB: 1QGT [1]) and SHBs (PDB: 8YMJ [2]) structures and on the preS protein sequence from genotype A are colored with positional score from the MULAN prediction. b. Residues in the capsid (PDB: 1QGT[1]) and SHBs (PDB: 8YMJ[2]) structures on the preS protein sequence from genotype C are colored with positional score from the MULAN prediction.

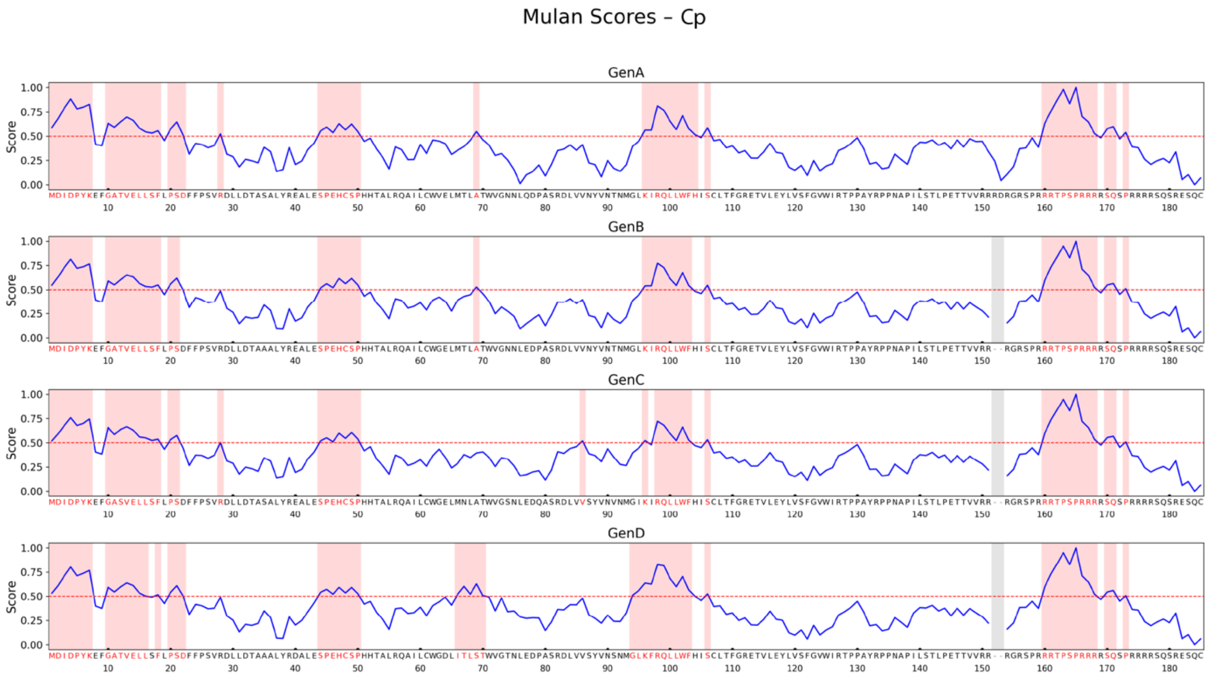

**Figure S2**. **MuLAN positional scores plotted for each genotype of Cp.** Predicted interface residues for Cp correspond to position 1-7, 10-18, 20-22, 28, 44-50, 69, 96-104, 106, 160-168, 170-171, 173 in the sequence for genotype A. Predicted interface residues for Cp correspond to position 1-7, 10-18, 20-21, 44-50, 69, 96-103, 106, 160-168, 170-171, 173 in the sequence for genotype B. Predicted interface residues for Cp correspond to position 1-7, 10-18, 20-21, 28, 44-50, 86, 96, 98-103, 106, 160-168, 170-171, 173 in the sequence for genotype C. Predicted interface residues for Cp correspond to position 1-7, 10-16, 18, 20-22, 44-50, 66-69, 96-103, 106, 160-168, 170-171, 173 in the sequence for genotype D. The red background and red characters highlight MuLAN [3] scores above 0.5.

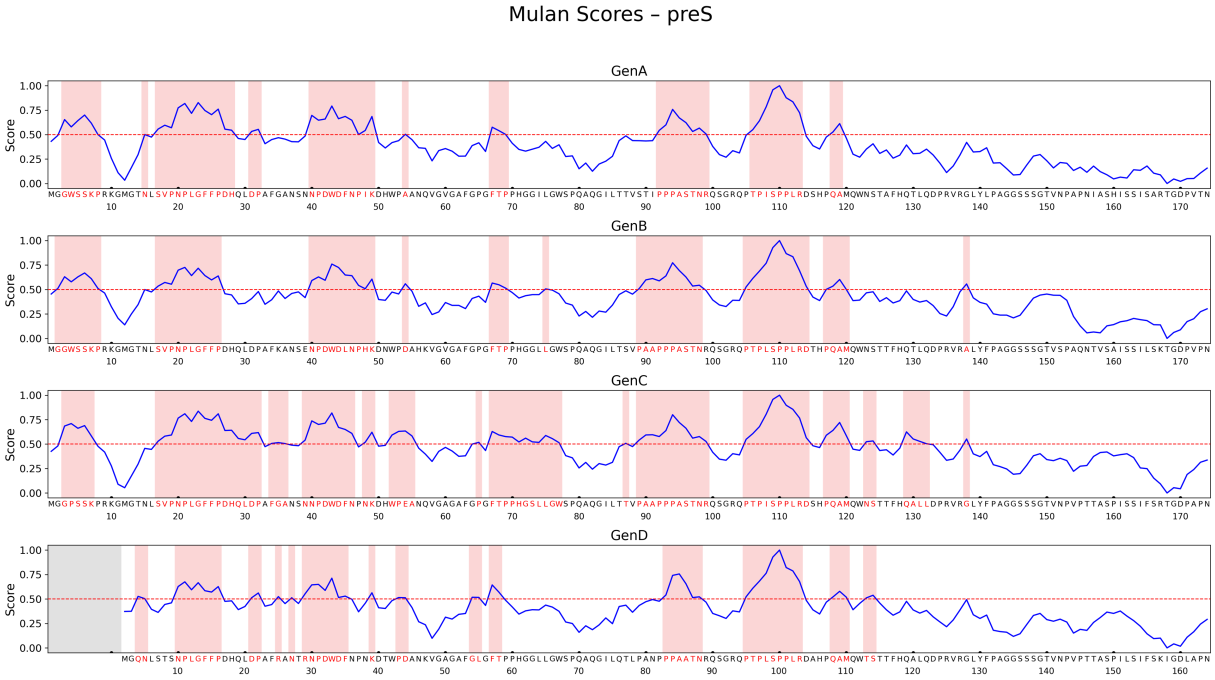

**Figure S3**. **MuLAN positional scores plotted for each genotype of preS.** Predicted interface residues for preS correspond to position 3-8, 15, 17-28, 31-32, 40-49, 54, 67-69, 92-99, 106-113, 118-119 in the sequence for genotype A. Predicted interface residues for preS correspond to position 2-8, 17-26, 40-49, 54, 67-69, 75, 89-98, 105-114, 117-120, 138 in the sequence for genotype B. Predicted interface residues for preS correspond to position 3-7, 17-32, 34-36, 39-46, 48-49, 52-55, 65, 67-77, 87, 89-99, 105-114, 117-120, 123-124, 129-132, 138 in the sequence for genotype C. Predicted interface residues for preS correspond to position 3, 4, 10-16, 21,22,25,27,29-35,39,43,44,54,55,57,58,83-88,95-103,108-110,113,114 in the sequence for genotype D. The red background and red characters highlight MuLAN [3] scores above 0.5.

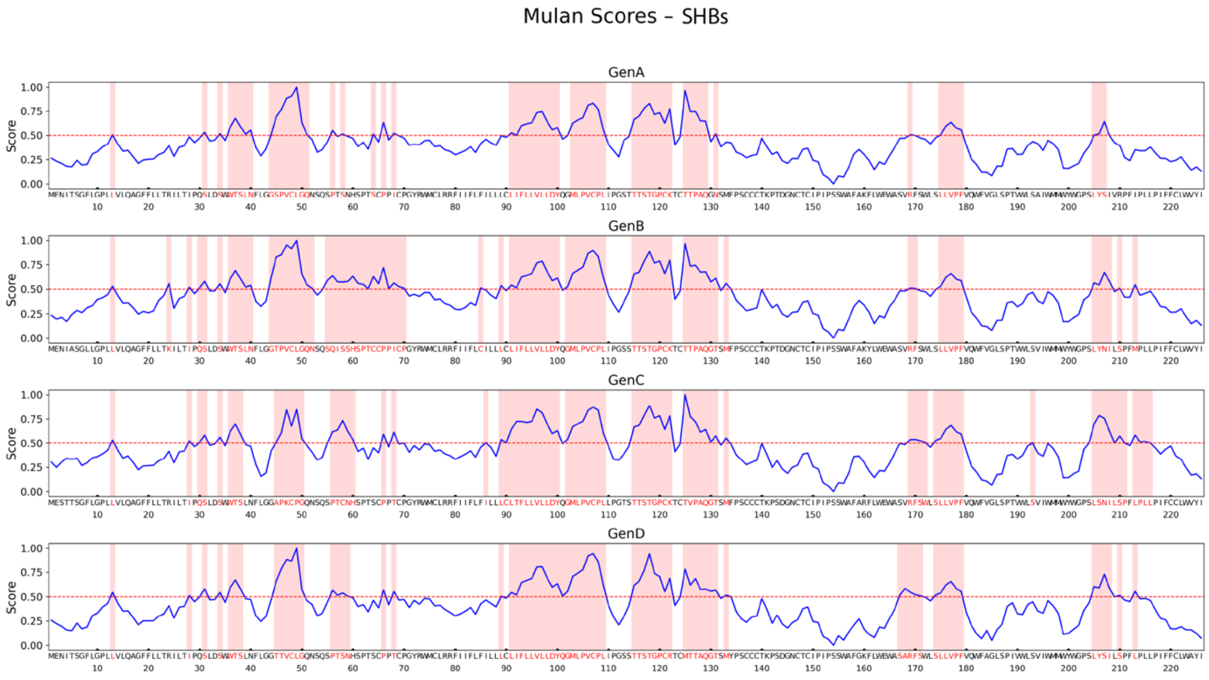

**Figure S4. MuLAN positional scores (left axis) plotted for each genotype of SHBs.** Predicted interface residues for SHBs correspond to position 13, 31,34,36-40, 44-51, 56, 58, 64, 66, 68, 91-100, 103-109, 115-122, 125-129, 131, 169, 175-179, 205-207 in the sequence for genotype A. Predicted interface residues for SHBs correspond to position 13, 24, 28, 30-31, 34, 36-40, 44-52, 55-70, 85, 89, 91-100, 102-109, 115-122, 125-131, 133, 169-170, 175-179, 205-208, 210, 213 in the sequence for genotype B. Predicted interface residues for SHBs correspond to position 13, 28, 30-31, 34, 36-38, 45-50, 56-60, 66, 68, 89-100, 102-109, 115-122, 125-131, 133, 169-172, 174-179, 193, 205-211, 213-216 in the sequence for genotype C. Predicted interface residues for SHBs correspond to position 13, 28, 31, 34, 36-38, 45-50, 56-59, 66, 68, 91-109, 115-122, 125-131, 133, 169-172, 174-179, 193, 205-208, 210, 213 in the sequence for genotype D. The red background and red characters highlight MuLAN [3] scores above 0.5.

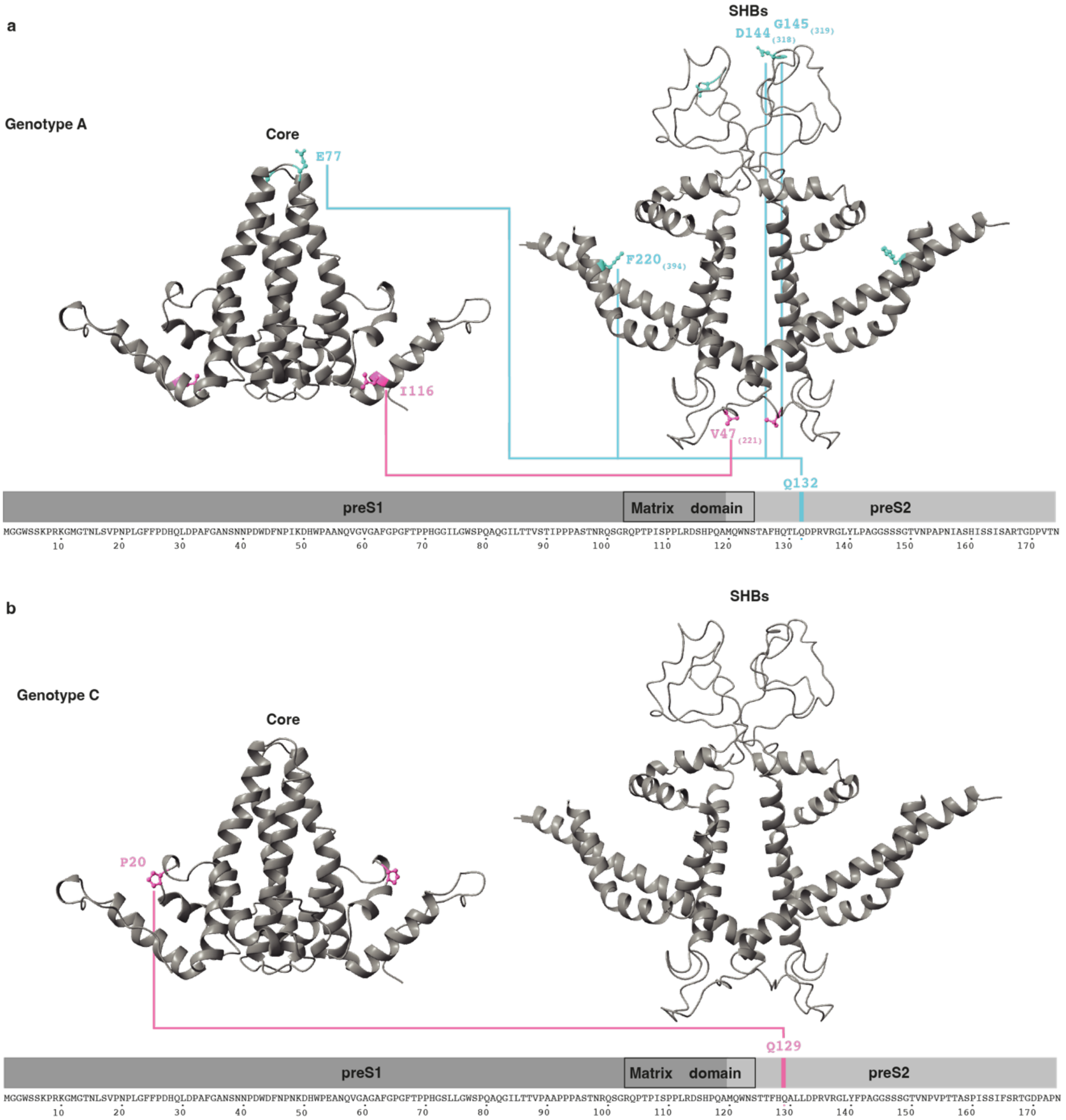

**Figure S5**. **Visualization of co-evolution clusters on the Cp dimer, preS and SHBs.** (a) Genotype A shows two co-evolution events. Amino-acid numbers on SHBs structure are given for SHBs numbering and in brackets for LHBs. (b) Genotype C shows one co-evolution event. The residue positions of each cluster are connected by lines. Structures for the plots are from Cp (1QGT.pdb [1] ) and SHBs (PDB: 8YMJ [2]) and the sequence is given for the preS domain.

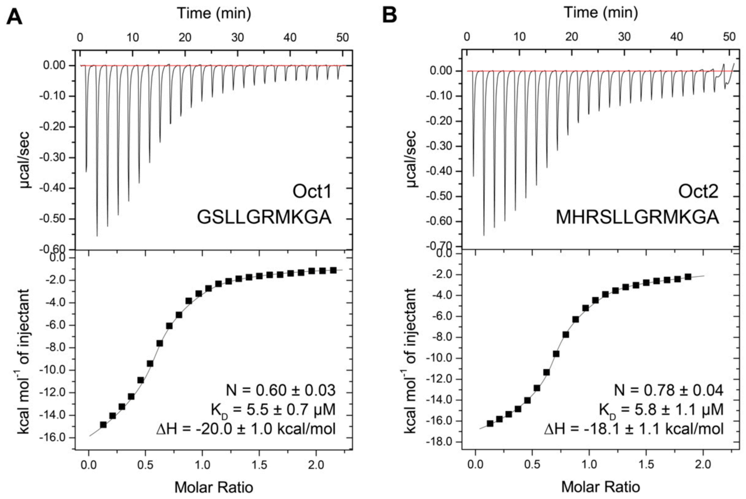

**Figure S6. ITC curves showing the interaction between Cp149 capsids and Oct peptides.** A) ITC raw data and binding isotherms displaying the titration of Oct1 to Cp149 reassembled capsids. B) ITC raw data and binding isotherms for the titration of Oct2 to Cp149 reassembled capsids. Experiments were done in 50 mM HEPES buffer pH 7.5 at 25 °C, with [Cp149monomer] = 40 µM and [Oct] = 400 µM. Sequences of both peptides are shown, as well as the resulting thermodynamic parameters.

**Table S1. NMR experimental details of solid-state NMR measurements.** Sw stands for spectral width. Experiments were recorded on an 800 MHz wide-bore spectrometer at a 17.5 kHz MAS frequency and at a sample temperature estimated at 4 °C. The $^{15}$N-insert was removed from the probe to increase the signal-to-noise ratio.

| **Sample** | **Cp149+Oct1** | **Cp149+Oct2** |
|---|---|---|
| Experiment | 2D DARR | 2D DARR |
| **Transfer 1** | HC-CP | HC-CP |
| Field [kHz] | 67.0 ($^{1}$H)<br>50 ($^{13}$C) | 65.7 ($^{1}$H)<br>50 ($^{13}$C) |
| Shape | Tangent $^{1}$H | Tangent $^{1}$H |
| $^{13}$C carrier [ppm] | 58.6 | 58.6 |
| time [ms] | 1.0 | 0.8 |
| **Transfer 2** | DARR | DARR |
| Field [kHz] | 17.5 ($^{1}$H) | 17.5 ($^{1}$H) |
| $^{13}$C carrier [ppm] | 100 | 100 |
| time [ms] | 20 | 20 |
| $t_1$ increments | 2560 | 2560 |
| sw ($t_1$) [kHz] | 93.75 | 93.75 |
| Acq. time ($t_1$) [ms] | 13.7 | 13.7 |
| $t_2$ increments | 3072 | 3072 |
| sw ($t_2$) [kHz] | 93.75 | 93.75 |
| Acq. time ($t_2$) [ms] | 16.4 | 16.4 |
| $^{1}$H decoupling | SPINAL64 | SPINAL64 |
| Field [kHz] | 90 | 90 |
| Interscan delay d1 [s] | 2.6 | 2.6 |
| Number of scans | 12 | 16 |
| Measurement time | 22 h 36 | 1 d 6 h |
